# The Dark Side of Light: Adolescent Light Exposure at Night Disrupts Hippocampal Clocks and Plasticity-Related Signaling in a Sex-Dependent Manner

**DOI:** 10.64898/2026.11.23.768669

**Authors:** Ashita Bhan, Janhvi Kartik, Pablo Bonilla, Alessandra Porcu

## Abstract

In today’s society, adolescents are being increasingly exposed to artificial blue light at night from electronic devices, raising concerns about its effects on brain development and mental health. The hippocampus, and specifically the dentate gyrus (DG) regulates memory formation and itself operates as a subordinate circadian oscillator detecting changes in the light environment. However, whether light at night during adolescence can impair hippocampal clock and function remains unknown. To begin exploring this, we exposed adolescent mice to a light cycle disruption (LCD) paradigm consisting of 12hrs of light followed by 5hrs of darkness and 7hrs of blue-enriched light for 5 consecutive days per week, followed by a standard 12 hours of light and 12 hours of darkness (12L:12D) for 2 days, over a 4-week period. Control animals were maintained under standard 12L:12D. Female and male mice were then tested for novel object recognition (NOR) and novel object location (NOL), and brain tissues were collected for clock gene expression, neuroplasticity markers, and dendritic spine analyses. We found that male mice exposed to LCD spent less time exploring novel objects and locations, and more time with familiar ones, a pattern consistent with impaired recognition and spatial memory, while female mice showed preserved memory performance despite LCD exposure. LCD-exposed male mice showed altered hippocampal daily expression rhythms of *Clock* and CRY2 levels compared to male control and LCD-exposed females. In addition, we found that LCD-exposed males exhibited reduced p-CREB expression along with reduced *Bdnf* and dendrite spines numbers in the DG. Interestingly, returning LCD-exposed males to standard 12L:12D conditions for 4 weeks were sufficient to reverse both the behavioral and molecular phenotypes. Together, these results suggest that chronic exposure to light at night during adolescence impairs memory in male mice, through alteration in hippocampal circadian clock regulation and plasticity-related mechanisms within the DG.

## Introduction

Adolescence is a developmentally sensitive period during which the hippocampus, particularly the dentate gyrus (DG), continues to mature and remains especially susceptible to change in the environments (Zehr et al. 2008; Shome et al. 2018). Today, approximately 80% of the world’s population lives under light-polluted night skies (Falchi et al. 2016), and several studies in humans have shown that exposure to light pollution is associated with increased mental disorders, particularly among adolescents (Menculini et al. 2024; Sharma et al. 2025; Paksarian et al. 2020). Furthermore, a recent longitudinal human study has shown that higher state average nighttime light intensity is associated with higher state Alzheimer’s Disease prevalence (Voigt et al. 2024). In modern society, adolescents are increasingly exposed to artificial light at night due to several light-emitting electronic devices and sleep schedules regulated by school time and social demands (Guindon et al. 2024; Ricketts et al. 2022). Furthermore, dim light at night has been shown to impair cognition (Fonken et al. 2012). Therefore, understanding how alteration in the light environment influences cognitive function during adolescence is essential for elucidating the potential consequences of light exposure on brain maturation and memory.

Light transmits information to the brain via image forming cells (rods and cones) and via intrinsically photosensitive retinal ganglion cells (ipRGCs) (Schmidt et al. 2011; Altimus et al. 2008). The ipRGCs represent a small subset of retinal ganglion cells that express melanopsin, a photopigment with peak sensitivity to blue light (~480 nm) (Brainard et al. 2001; Berson et al. 2002). IpRGCs play a critical role in circadian photoentrainment by transmitting environmental light information directly to the suprachiasmatic nucleus (SCN), the master circadian clock, via the retinohypothalamic tract (RHT) (Kofuji et al. 2016). Upon light stimulation, ipRGCs release glutamate onto the SCN, activating glutamatergic receptor and promoting Ca^2+^ influx. This triggers downstream signaling cascades that regulate the expression of core clock genes and thereby synchronizing the endogenous circadian clock with the external light–dark cycle (Partch et al. 2014; Hattar et al. 2002; Berson et al. 2002; Hannibal 2002). Importantly, ipRGCs project to several parts of the brain beyond the SCN, including the amygdala, and the lateral habenula, thereby influencing a wide range of brain functions (Güler et al. 2008). However, direct ipRGC projections to brain regions critically involved in learning and memory, such as the hippocampus, have not been identified, suggesting that light may influence cognitive functions through indirect neural pathways (LeGates et al. 2012; Fernandez et al. 2018a).

The hippocampus contains an intrinsic circadian clock characterized by rhythmic expression of clock genes. Although these local molecular oscillations can persist independently of direct SCN input, their phase and synchronization are influenced by the central circadian system (Wang et al. 2009; Harbour et al. 2014; Walters et al. 2026). Previous studies have shown that clock genes in DG granule cells gate time-of-day neuronal excitability and memory processing and selective knockdown of Bmal1 decreases cognition (Gonzalez et al. 2023). Moreover, studies have shown that long day exposure enhanced long-term recognition memory, and this effect was abolished by loss of the melanopsin (Dellapolla et al. 2017), In addition, *Per1* rhythmic expression play a critical role in hippocampal memory formation (Bellfy et al. 2023), suggesting that maintaining circadian integrity within the DG might be critical for memory function. Furthermore, our previous work has shown that exposure to chronic light cycle disruption during adolescence reduced recognition memory along with altered daily *Per1* expression and neuron activity in the DG (Bonilla et al. 2024).

Beyond the regulation of core clock genes, light exposure and circadian rhythms can also influence molecular pathways involved in synaptic plasticity and memory formation (Siraji et al. 2023; Hartsock and Spencer 2020; Gerstner and Yin 2010). Among these, the cAMP response element-binding protein - Brain Derived Neurotropic Factor (CREB-BDNF) signaling axis plays a critical role in activity-dependent gene expression and synaptic plasticity and have been shown to be affected by light exposure (Tao et al. 1998; Esvald et al. 2020)(Lu et al. 2008; Ortega-Martínez 2015; Dellapolla et al. 2017). Similarly, CaMKv, a calcium/calmodulin-binding pseudokinase, contributes to dendritic spine maintenance and spatial memory (Liang et al. 2016). Together, these findings suggest that altered light exposure may influence hippocampal memory through changes in both circadian regulation and plasticity-related mechanisms. Here, we investigated whether chronic exposure to altered light conditions during adolescence affects hippocampal clock gene expression, CREB–BDNF signaling, CaMKv expression, and dendritic spine density within the DG, and whether these effects are reversible.

## Results

### Exposure to chronic light at night during adolescence impaired hippocampal memory exclusively in male mice

To investigate the effects of altered light environments on DG-dependent memory during adolescence, we exposed adolescent mice (postnatal day 30) to light cycle disruption (LCD) paradigm or control conditions (Bonilla et al. 2024). Following four weeks of exposure (postnatal day 60), memory tasks were conducted at ZT8-ZT9 using the novel object recognition (NOR) and novel object location (NOL) tests, two well-established tests for assessing dentate gyrus (DG) dependent recognition memory and spatial memory, respectively (Jessberger et al. 2009; Lee et al. 2005). ZT8-9 was selected based on previous findings demonstrating peak memory performance during this time window (Bellfy et al. 2023). In the NOR task (Fig. 1A-C), LCD-exposed male mice showed a significant reduction in time spent exploring the novel object (Fig. 1D), increased time exploring the familiar object (Fig. 1E), and a significantly decreased discrimination index compared to male controls (Fig. 1F), indicating impaired recognition memory. In contrast, LCD-exposed females showed no significant differences compared to controls in time spent exploring the novel object and discrimination index (Fig. 1D & F). However, LCD-exposed female mice spent significantly more time investigating the familiar object (Fig. 1E).

**Figure 1:**
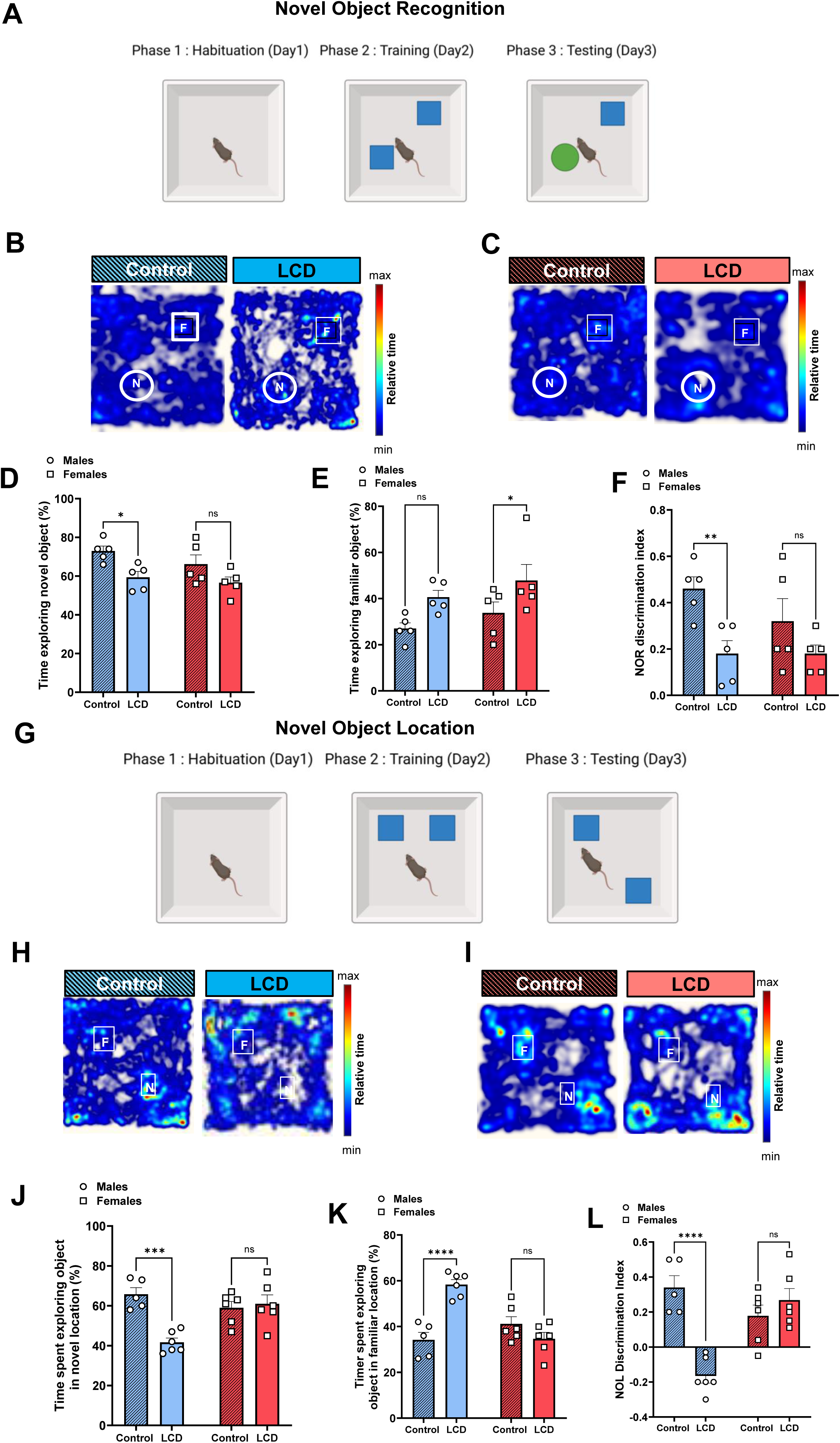
LCD exposure induces hippocampal memory deficits exclusively in male mice. Effects of LCD on Novel Object Recognition (A) and Location (G): For NOR, bar graphs show the percentage of time spent exploring the novel object (D), the familiar object (E), and the discrimination index (F) in male and female mice. For NOL, bar graphs show the percentage of time spent exploring the familiar object location (K), the novel object location (J), and the discrimination index (L) in male mice. Panels (B, C, H, I) show representative heatmaps of mouse movement during the respective tasks. Individual data points represent individual mice. Data are presented as mean ± SEM, *p<0.5, **p < 0.01, ***p < 0.001, ***p < 0.0001, Two-way ANOVA.

In the NOL task, (Fig. 1G-I) LCD-exposed male mice showed significant reductions in time spent exploring the object in the novel location (Fig. 1J), increased time spent investigating the familiar object (Fig. 1K), and significantly decreased NOL discrimination index relative to controls (Fig. 1L), indicating a profound impairment in spatial memory. LCD-exposed females, however, showed no significant differences from controls across any NOL measure (Fig. 1J-L). Together, these findings demonstrate that LCD exposure during adolescence selectively impairs recognition and spatial memory in males, while leaving memory performance largely intact in females, revealing a sex-dependent vulnerability to light cycle disruption during adolescence.

### Exposure to chronic light at night during adolescence alters DG molecular clock but not *Bdnf* rhythms in a sex-dependent manner

To test whether memory impairments are associated with altered clock gene expression, we analyzed *Clock*, *Per1* and CRY2 daily expression patterns in the DG at ZT2, ZT8, ZT16, and ZT22. We found antiphase daily *Clock* expression patterns in LCD-exposed male mice compared to control, while no effect was observed in LCD-exposed female mice (Fig. 2A-D). No effect was observed in *Per1* daily expression patterns in both male and female mice after LCD exposure (Fig. 2E-H). Interestingly, we found that LCD-exposed male mice showed significantly reduced CRY2 expression at ZT16 compared to control (Fig. 2I-J) while LCD-exposed females had significantly higher levels of CRY2 across all examined ZT time points compared to control (Fig.2K-L). Given that BDNF levels oscillate across tissues with distinct phasing in responses to clock perturbations (Ehrhardt et al. 2024) and light exposure (Dellapolla et al. 2017), we analyzed *Bdnf* daily expression in the DG at ZT2, ZT8, ZT16, and ZT22. We found no significant difference in both male and female mice after LCD exposure compared to control (Fig.3A-D). These findings suggest that LCD selectively alters the rhythmic expression of specific genes, rather than broadly disrupting hippocampal molecular rhythms.

**Figure 2:**
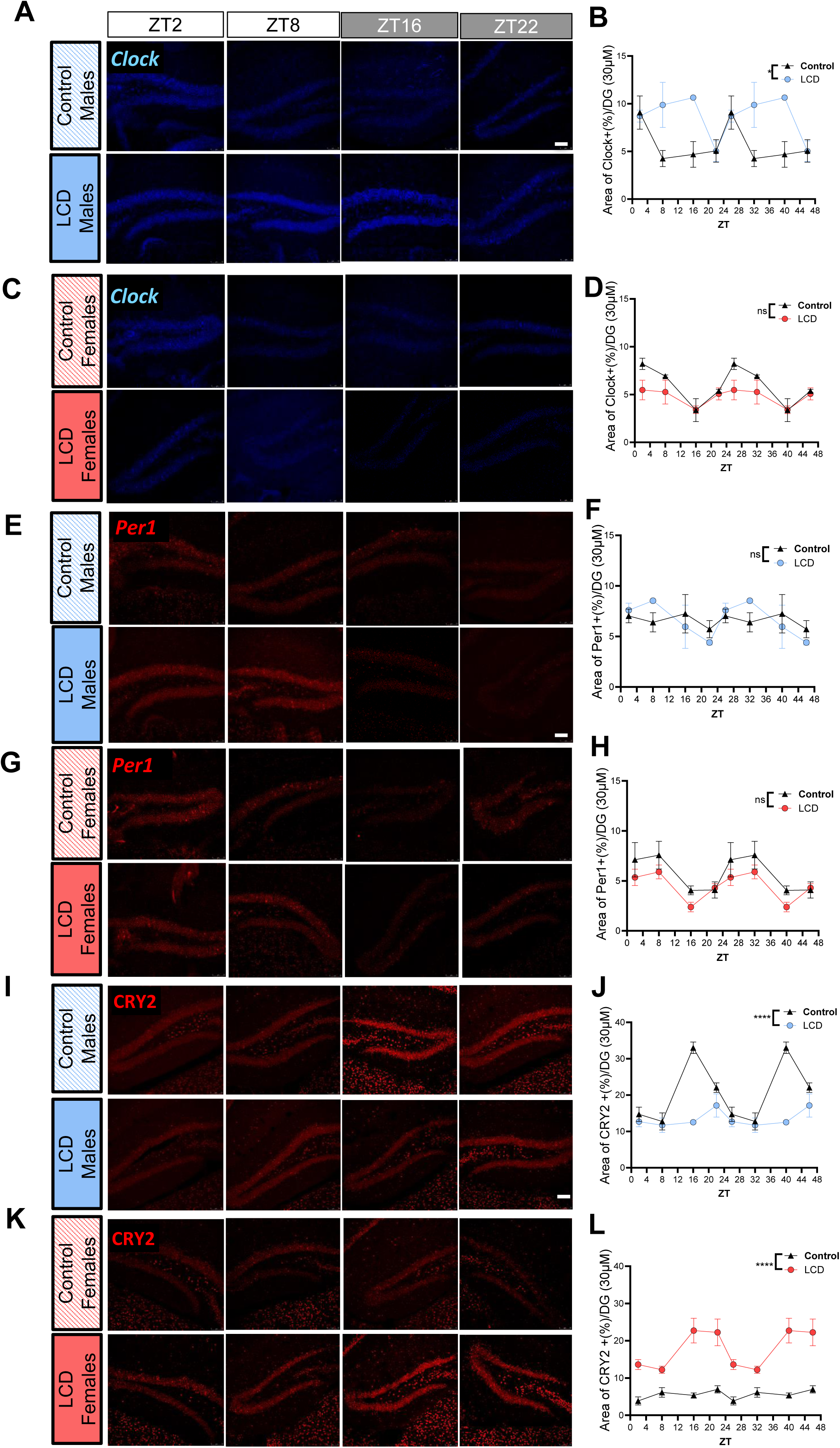
LCD exposure alters hippocampal molecular clock genes expression in a sex-specific manner. Representative confocal micrographs showing *Clock* (blue,A & C), *Per1* (red, E & G) and CRY2 (red, I & K) in Control and LCD; males and females (Scale bar, 75 μm). Line graphs depict double-plotted rhythms of *Clock* (B & D), *Per1* (F & H) and CRY2 (J & L) across four circadian timepoints (ZT2, ZT8, ZT16, and ZT22). Data are double plotted to facilitate visualization of rhythmicity. *Clock* and *Per1* mRNA expressions were quantified by fluorescence in (ISH-RNAscope). CRY2 protein expression was assessed by immunofluorescence (IF). Individual data points represent individual mice. All data are presented as mean ± SEM; *p<0.5, **p < 0.01 ****p < 0.0001, ns= no significance. Two-way ANOVA.

**Figure 3:**
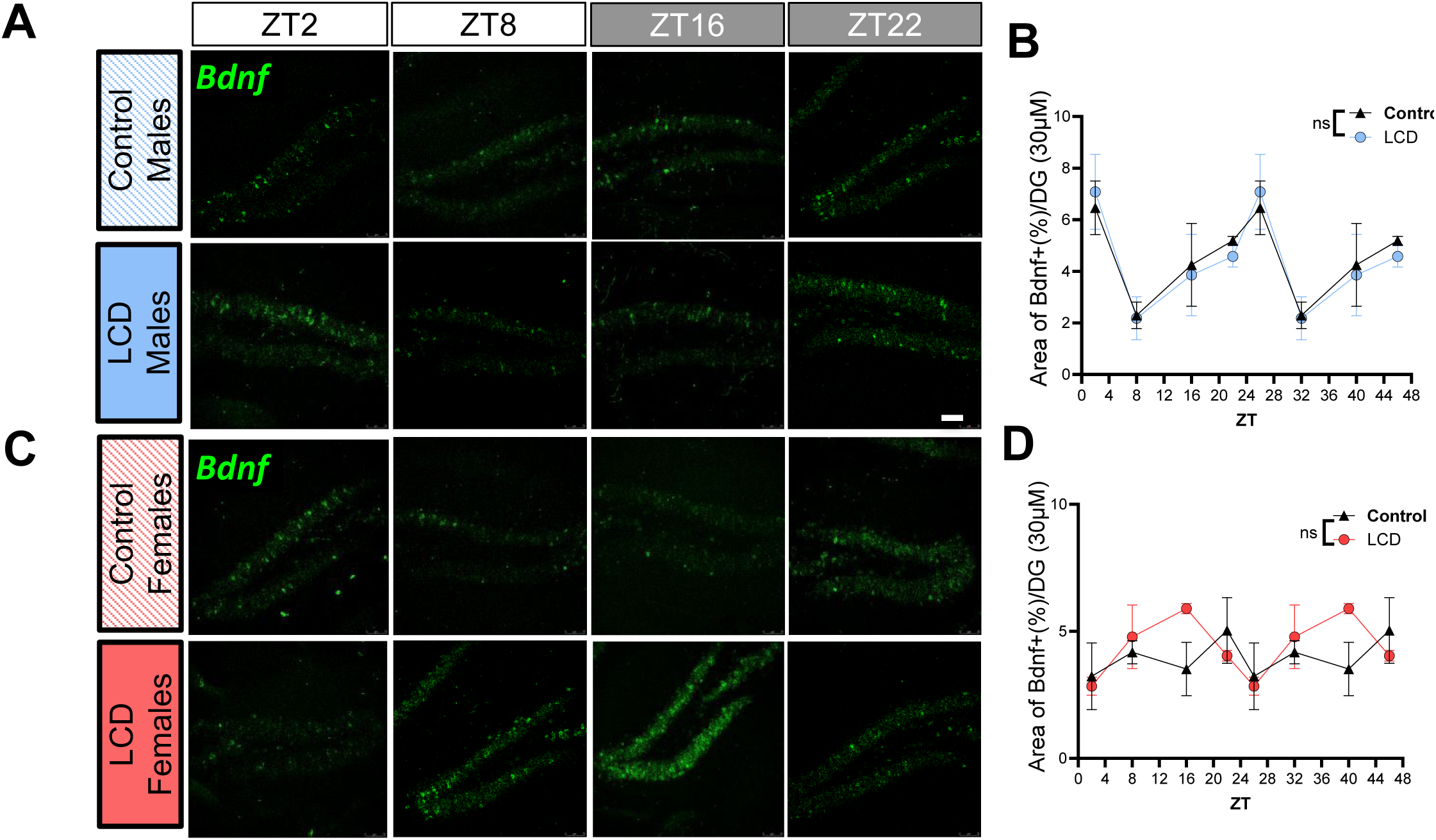
LCD exposure has no effect on Bdnf daily expression in male and female mice. Representative confocal micrographs showing *Bdnf (*green, A & C) in Control and LCD (Scale bar, 75 μm); males and females. Line graphs depict double-plotted rhythms of *Bdnf* in males (B) and females (D). Individual data points represent individual mice. All data are presented as mean ± SEM; ns= no significance. Two-way ANOVA.

### Exposure to chronic light at night during adolescence reduces hippocampal plasticity-related pathways and dendritic integrity exclusively in male mice

To investigate the molecular mechanism underlying the sex-dependent memory deficits observed in NOR and NOL tasks, we examined the expression of pCREB, BDNF, CaMKV and dendritic density in the DG, 45 minutes following behavioral testing, when activity-dependent transcriptional responses are expected to be elevated (Guzowski et al. 2001; Wu et al. 2001), using immunofluorescence, RNAscope in situ hybridization and Golgi-Cox staining (Fig. 4A). pCREB immunofluorescence, revealed a significant reduction in pCREB+ cell density within the DG of LCD-exposed males compared to controls (Fig. 4B, C). This decrease was absent in LCD-exposed females, whose pCREB+ cell density showed no difference compared to controls (Fig. 4B, C), suggesting that the impairment of upstream transcriptional activation is sex-specific. Consistent with a impaired pCREB dependent pathway, RNAscope in situ hybridization revealed a significant reduction of *Bdnf* mRNA expression in LCD-exposed male mice relative to controls (Fig. 4D, E), while no difference was observed in LCD-exposed females (Fig. 4D, E), further reinforcing the male-specific nature of these molecular alterations. Given that CaMKV has been identified as an activity-dependent pseudokinase required for the maintenance of dendritic spine density and spatial memory (Liang et al. 2016), we next examined its expression after LCD and memory tasks. RNAscope in situ hybridization revealed a significant reduction of *Camkv* mRNA expression in LCD-exposed males, while females remained unaffected (Fig. 4F, G). Consistent with the reduced *Camkv* expression, Golgi-Cox analysis of DG dendrites revealed a significant reduction in spine density per unit dendritic length in LCD-exposed male mice compared to control (Fig. 4H, I), suggesting that these molecular alterations are accompanied by structural changes in synaptic architecture.

**Figure 4:**
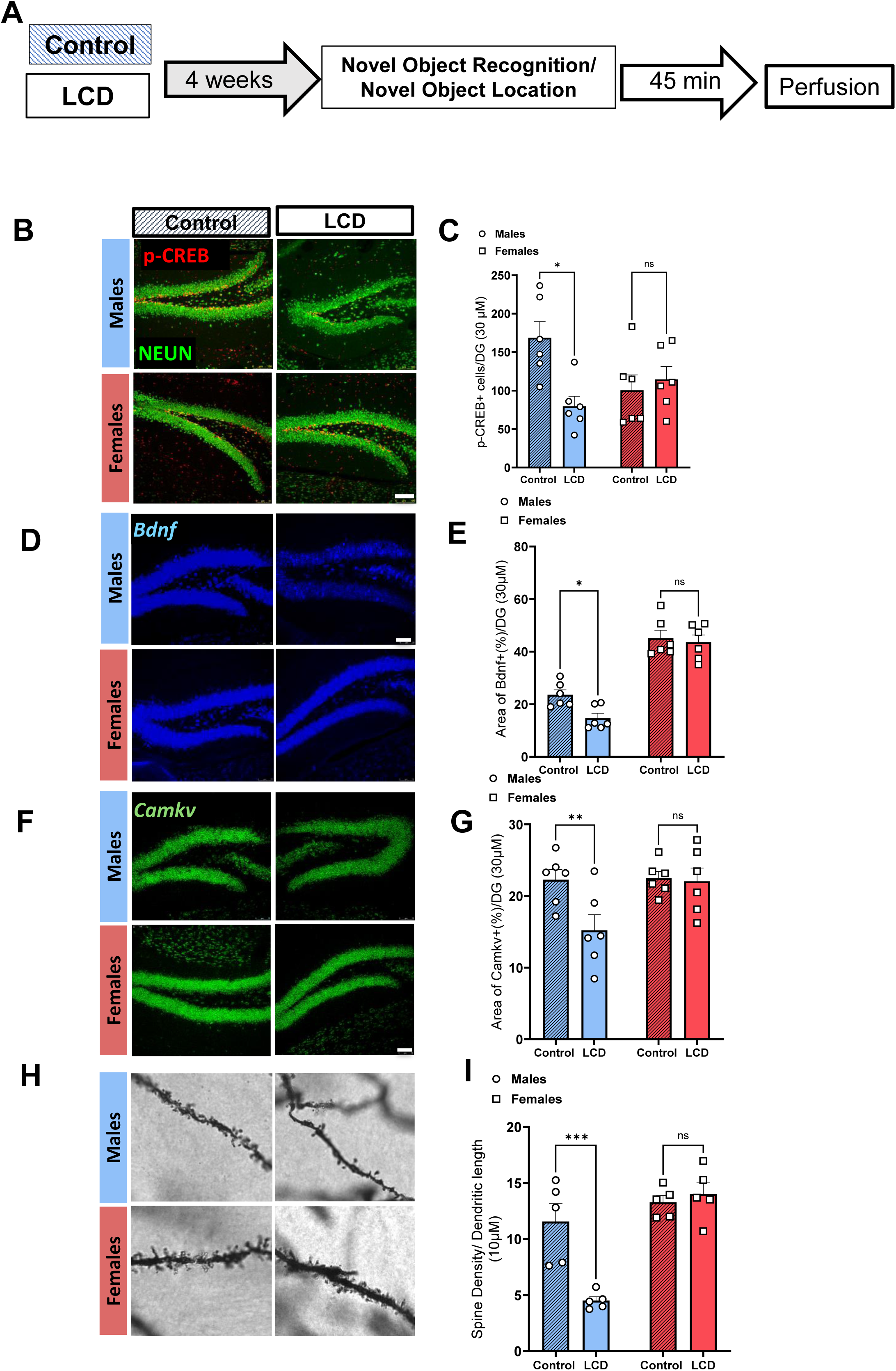
LCD exposure reduces hippocampal plasticity-related pathways and dendritic integrity in a sex-specific manner. Schematic of experimental design after 4 weeks of exposure to LCD, mice undergo memory tasks and after are perfused for molecular analysis (A). Representative confocal micrographs showing (B) phosphorylated cAMP-responsive element binding protein (p-CREB) (red) protein expression detected by IF in control and LCD in male and female mice. (Scale bar 100μm). Bar graphs show quantification of p-CREB in the DG following LCD or control exposure in males & females, respectively (C). Representative confocal micrographs *Bdnf* (blue) and *Camkv* (green) (D & F) show mRNA expression detected by ISH-RNAscope in the DG of control and LCD-exposed male and female mice (Scale bar, 75 μm). Bar graphs show quantification of *Bdnf* (E) in males and females respectively, a key neurotrophic factor regulating synaptic plasticity and memory formation. While bar graphs show quantification of *Camkv* (G) in males and females respectively, which is essential for calcium-dependent signaling and memory consolidation. Representative confocal micrographs showing spine density via Golgi staining in control and LCD in (H) male and female mice (Scale bar 8μm). Bar graphs show spine density / 10um of dendritic branch length in (I) in males and females respectively in the DG. Bar graphs show quantification. Individual data points represent independent mice. Data are shown as mean ± SEM.*p<0.5, **p < 0.01, ***p < 0.001. Two-way ANOVA.

### Exposure to chronic light at night during adolescence does not affect neurogenesis or neuronal death in the DG

To test whether memory deficits are associated with altered neurogenesis or cell death we quantified Doublecortin (DBC) expression by immunofluorescence and assessed apoptotic cell death using the TUNEL assay. We found that the total number of DBC+ cells was unchanged in both male and female mice following LCD exposure (Fig. 5 A, B, C). Furthermore, TUNEL+ cell counts were negligible in both LCD-exposed males and females, with no significant difference compared with their respective controls, and remained significantly lower than those observed with DNase-treated positive control tissue (Fig. 6A–D).

**Figure 5:**
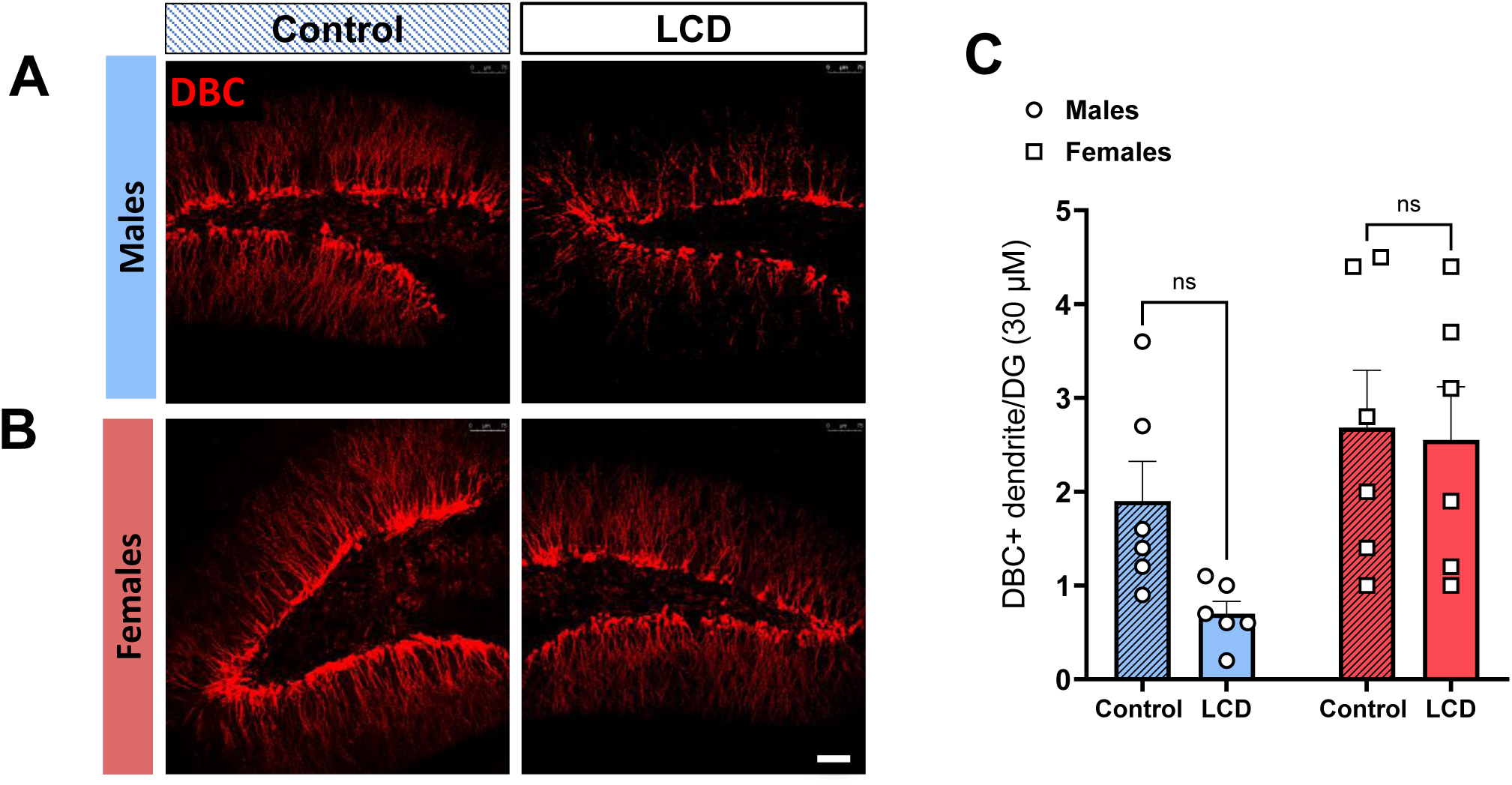
LCD exposure does not affect neurogenesis. Representative confocal micrographs showing DBC (red) protein expression detected by IF in control and LCD in (A) male mice and (B) female mice (Scale bar 75μm). Bar graphs show DBC expression (C) in males and females in the DG. Individual data points represent independent mice. Data are shown as mean ± SEM; ns= no significance. Two-way ANOVA.

**Figure 6:**
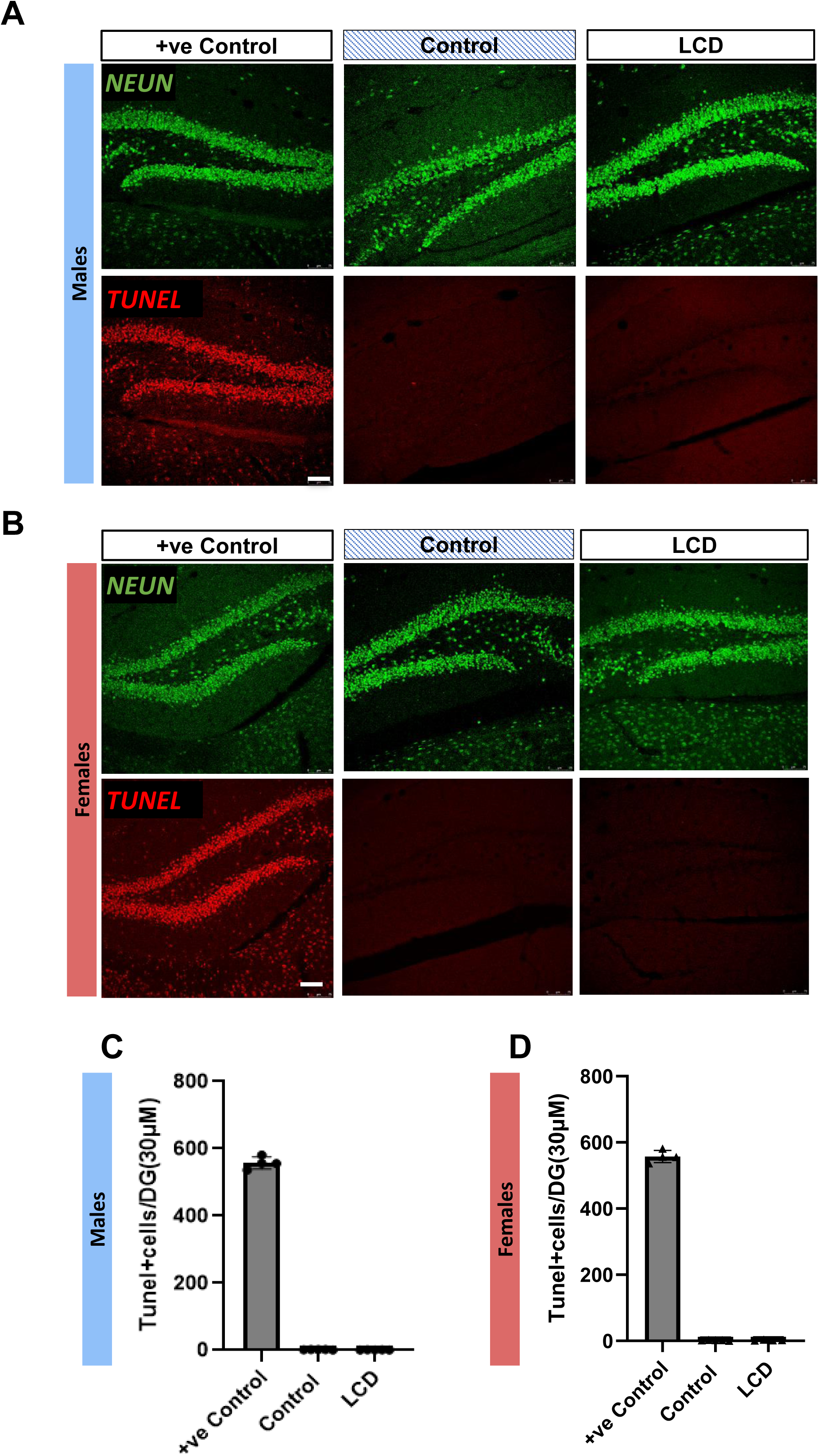
LCD exposure does not induce neuronal death. Representative confocal micrographs showing TUNEL+ (red) expression detected by TUNEL Assay in control and LCD in male mice (A) and in female mice (B) (Scale bar 75μm). For Positive control, the tissue was treated with DNase. Bar graphs show TUNEL assay quantification in males (C) and in females (D) in the DG. Individual data points represent independent mice. Data are shown as mean ± SEM; ns= no significance. Two-way ANOVA.

### Restoration to 12L:12D Light-Dark Cycle Reverses LCD-Induced Memory Deficits in Adult Male Mice

To determine whether the memory deficits induced by LCD exposure during adolescence persist into adulthood or can be reversed by restoring a normal light-dark cycle, we examined recognition and spatial memory in adult male mice following their return to a standard 12L:12D. To this aim, following the 4 weeks of LCD exposure or control condition, mice were kept under 12:12 light-dark conditions for four weeks (P60-P90), after which NOR and NOL performance and hippocampal plasticity related-markers were assessed (Fig. 7A).

**Figure 7:**
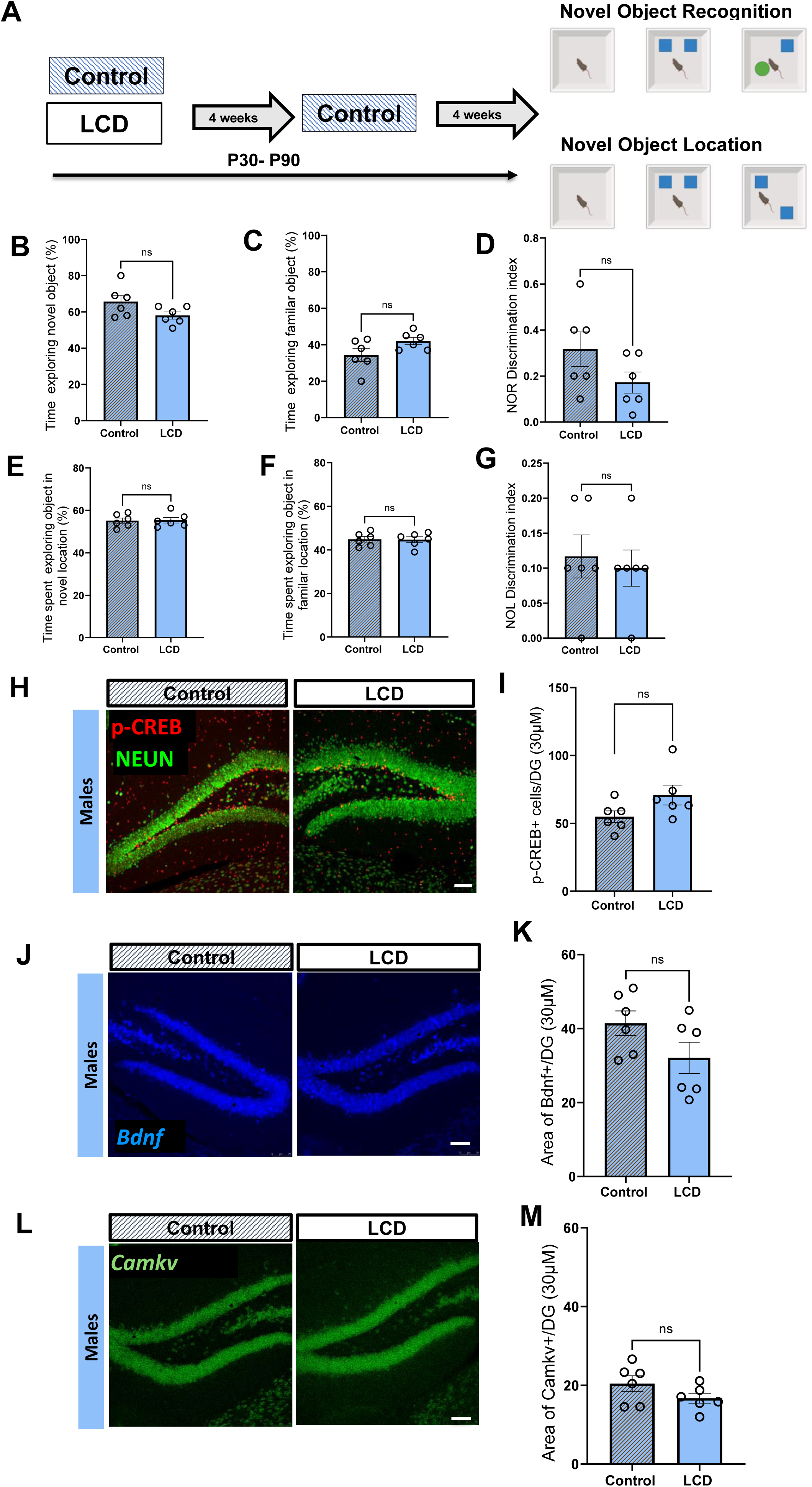
Restoration to 12L:12D light-dark cycle rescues LCD-induced memory deficits in adult male mice. Schematic reflecting following LCD exposure, male mice underwent a four-week washout under a 12:12 light–dark cycle and then tested for NOR and NOL (A). Bar graphs show the percentage of time spent exploring (B) the novel object, (C) the familiar object, and (D) the discrimination index in male mice. For NOL, bar graphs show the percentage of time spent exploring (E) the object in novel location, (F) the object in familiar location, and (G) the discrimination index in male mice. Furthermore, representative confocal micrographs showing p-CREB (red, H) protein expression detected by IF and *Bdnf* (blue, J) and *Camkv* (green, L) show mRNA expression detected by ISH-RNAscope in the DG in control and LCD (Scale bar 75μm). Bar graphs show quantification of p-CREB protein expression (I), and *Bdnf* and *Camkv* mRNA expression (K & M) was quantified in the DG of control and LCD-exposed male mice. Individual data points represent independent mice. Data are presented as mean ± SEM. ns= no significance Student’s t-test.

In the NOR task, LCD-exposed males that underwent 12L:12D restoration showed significant recovery of time spent exploring the novel and familiar object, and NOR discrimination index to levels comparable to controls (Fig. 7B-D). Spatial memory was similarly rescued, with 12L:12D restored male mice showing recovery of time spent exploring the object in the novel and familiar location, and NOL discrimination index compared to controls (Fig. 7E-G). Importantly, the behavioral rescue was accompanied by normalization of the hippocampal plasticity related cascade. No significant difference was observed in pCREB+ cell density in 12L:12D restored male mice compared to controls (Fig. 7H, I). RNAscope in situ hybridization also revealed no significant difference in *Bdnf* and *CamkV* expression in the DG between 12L:12D restored male mice and controls (Fig. 7J-M). Furthermore, 12L:12D restored females continued to show no memory impairment (Fig. 8A–F). Altogether, these behavioral findings establish that the cognitive phenotype induced by adolescent LCD in males is fully reversible following restoration of a standard light-dark cycle.

**Figure 8:**
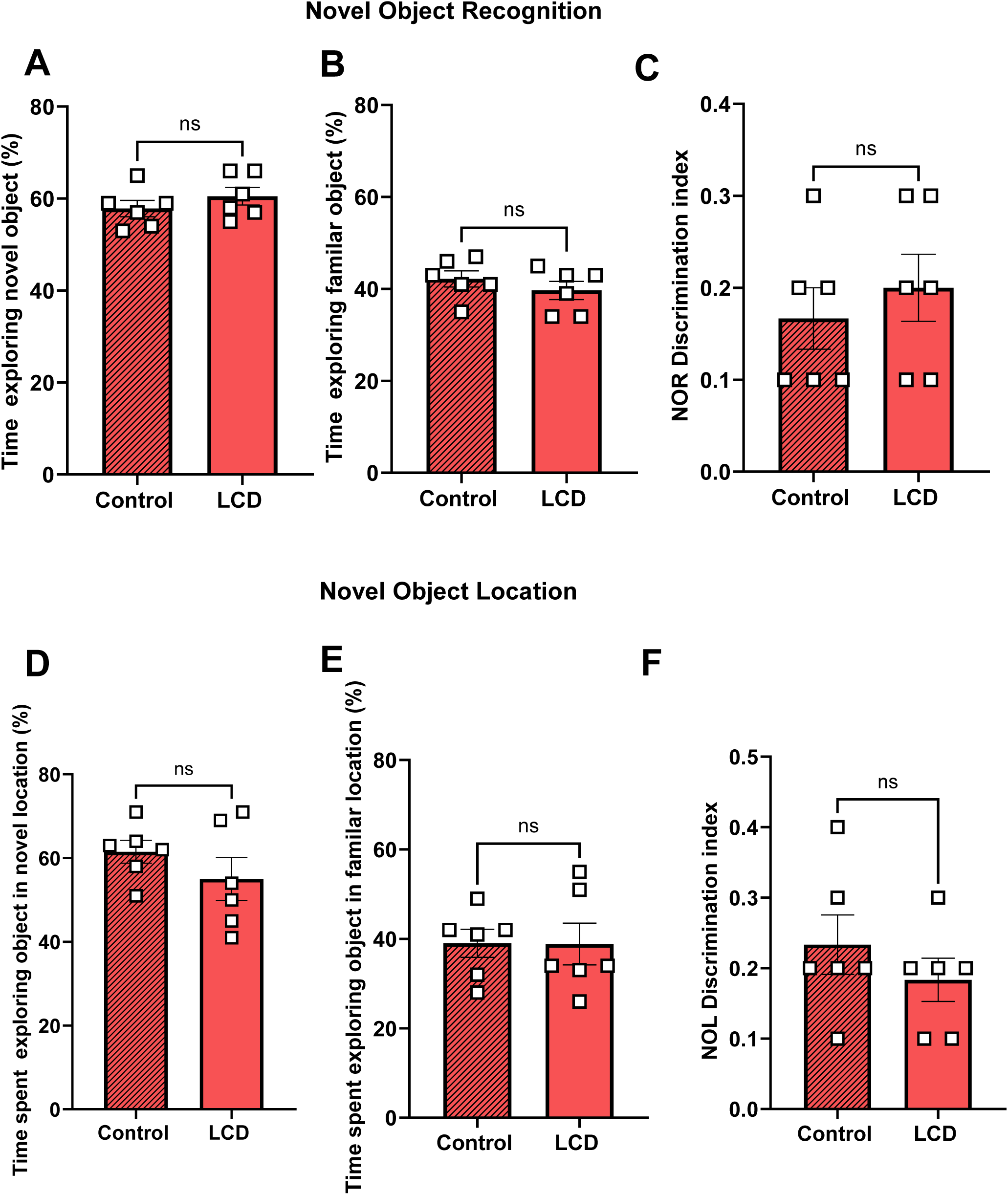
Restoration to 12L:12D Light-Dark in adult female mice. For NOR bar graphs show the percentage of time spent exploring the novel object (A), the familiar object (B), and the discrimination index (C) in female mice. For NOL, bar graphs show the percentage of time spent exploring the object in novel location (D), the object in familiar location (E), and the discrimination index in female mice (F). Individual data points represent independent mice. Data are presented as mean ± SEM. ns= no significance Student’s t-test.

## Discussion

Increasing exposure to artificial light at night during adolescence raises concerns about its potential impact on brain development and cognitive function. Although the hippocampus contains an intrinsic molecular clock and plays a critical role in memory formation (Smarr et al. 2014; Hasegawa et al. 2019), the consequences of adolescent nighttime light exposure on hippocampal circadian regulation and memory remain poorly understood. Here, we demonstrate that exposure to light at night during adolescence has lasting, sex-dependent effects on hippocampal molecular clock and memory function. We show that adolescent light cycle disruption selectively impairs recognition and spatial memory in male mice, accompanied by alterations in hippocampal molecular clock rhythms, plasticity-related signaling, and dendritic spine density. Importantly, these effects were reversible following restoration of a normal light-dark cycle, suggesting that the cognitive and molecular consequences of adolescent nighttime light exposure are not necessarily permanent. Together, these findings provide new evidence linking environmental light disruption during a critical period of brain development to hippocampal dysfunction and highlight sex-dependent differences in vulnerability and recovery.

The effects of artificial light at night have important implications, particularly during adolescence when the brain undergoes functional maturation (Arain et al. 2013). Human studies demonstrate that resynchronization of internal clocks is achievable after reduced evening light exposure (Wright et al. 2013). From a translational standpoint, this supports the view that modifiable light environments in adolescence can drive or reverse the phenotype. Adolescence is a period of ongoing DG maturation and heightened environmental sensitivity (Zehr et al. 2008; Shome et al. 2018), and epidemiological associations between nighttime light exposure, altered emotional and social behavior (Menculini et al. 2024; Sharma et al. 2025; Paksarian et al. 2020) make the persistence of long-term vulnerability a priority for future work.

An important finding of this current study is the prominent sex difference in susceptibility to altered light environments. Female mice exposed to LCD showed no memory impairment or molecular phenotypes that we observed in LCD-exposed male mice. However, LCD-exposed female mice spent significantly more time investigating the familiar object potentially reflecting altered exploratory behavior associated with the increased anxiety-like behavior previously observed in both male and female mice following LCD exposure in our earlier study (Bonilla et al. 2024). Moreover, in previous studies using chronic stress paradigms, male rats showed impairment in spatial and recognition memory compared to controls, whereas females exhibited faster recovery or even improved performance (Bowman et al. 2003; Conrad et al. 2003). This pattern suggests that male and female hippocampal circuits respond differently to sustained environmental challenges. However, the mechanisms underlying these sex-dependent differences remain to be elucidated. Furthermore, studies in humans have shown male subjects are more sensitive to that blue light compared to females (Chellappa et al. 2017). Sex-dependent differences in melanopsin expression within ipRGCs may contribute to this differential sensitivity, potentially influencing how light signals are transmitted to other brain regions.(Miguel et al. 2025).

Our study provides an in-depth investigation of the hippocampal clock which expands on our lab’s previous work (Bonilla et al. 2024). Here, we show that the effect of LCD on the local clock in the DG differs between males and females. LCD-exposed male mice showed reduced CRY2 rhythm in the DG. Meanwhile LCD-exposed females had upregulated daily expression of CRY2 compared to control. These contrasting responses may reflect sex-dependent differences in the sensitivity of the hippocampal molecular clock to environmental light disruption. Given the established influence of sex hormones on circadian regulation and hippocampal function, hormonal modulation of local clock machinery may contribute to these divergent responses (Goode et al. 2022; Yagi and Galea 2019). Such differences could influence the stability of memory-related molecular pathways under altered light conditions, potentially contributing to the preserved cognitive performance observed in females. Further studies should address whether LCD exposure during adolescence alters sex hormones levels, such as estradiol and testosterone.

In addition to alterations in molecular clock in the DG, our data also suggested that LCD impairs activity-dependent and not baseline plasticity signaling. Daily *Bdnf* expression did not differ between LCD-exposed mice compared to control. In contrast, Bdnf was reduced in LCD-exposed male mice 45 minutes after behavioral testing. Moreover, the reduced expression of pCREB+ cell following behavior testing is consistent with these findings, given the established role of phosphorylated CREB in regulating activity-dependent Bdnf transcription (Lu et al. 2008; Esvald et al. 2020). Moreover, CREB activity is required for memory formation and activity-dependent spine growth (Murphy and Segal 1997; Silva et al. 1998). BDNF too is known to support spine formation and persistence of memory (Tyler and Pozzo-Miller 2003; Bekinschtein et al. 2008). Similarly, CaMKv, a calcium/calmodulin-binding pseudokinase is known to contribute to dendritic spine maintenance and spatial memory (Liang et al. 2016). Thus, a reduction in pCREB, *Bdnf* and *Camkv* in the DG supports the memory deficits observed in LCD-exposed males. Conversely, the preservation of these plasticity-related markers in LCD-exposed females may underlie their maintained memory performance despite altered light exposure.

Across a wide range of stress paradigms, studies report that hippocampal-dependent learning is impaired. Chronic restraint stress causes atrophy of apical dendrites in pyramidal neurons (Watanabe et al. 1992; Magariños and McEwen 1995) and impairs spatial memory (V. Luine et al. 1994; Kleen et al. 2006). Acute stress can also reduce hippocampal spine density and produce memory deficits (Y. Chen et al. 2010). Stress during the juvenile and peripubertal period also alters hippocampal and cognitive function with effects that can persist into adulthood (Isgor et al. 2004). The effects of stress on hippocampal structure and memory are frequently sex-dependent. Females show less CA3 atrophy after chronic restraint than males (Galea et al. 1997), and stress can spare or even enhance spatial memory in females (Bowman et al. 2001; V. N. Luine et al. 2007). Together, these findings suggest that distinct environmental stressors may converge on common mechanisms affecting hippocampal dendritic integrity and memory, with males potentially exhibiting greater vulnerability than females.

Lastly, the relationship between environmental light exposure and hippocampal function is particularly interesting given that the hippocampus receives no direct projections from the SCN or ipRGCs (LeGates et al. 2012; Porcu et al. 2018), yet it contains its own clock that regulates DG-dependent processes such as learning and memory (Wang et al. 2009; Harbour et al. 2014; Walters et al. 2026). It is also known that a subset of ipRGCs projecting to the SCN can drive light-induced hippocampal learning, but their exact mechanism remains unknown (Fernandez et al. 2018b). Glucocorticoid signaling is a particularly feasible hypothesis given the abundance of glucocorticoid receptors in the hippocampus and their known interaction with CREB-dependent transcription (Reul and de Kloet 1985; Trollope et al. 2012; van Weert et al. 2017) and glucocorticoids also directly regulate spine formation (Liston and Gan 2011). Furthermore, light regulates the circadian expression of corticosterone and Glucocorticoid receptors in the DG (Coleman et al. 2016). Thus, alterations in glucocorticoid signaling may represent an additional mechanism through which chronic nighttime blue light exposure influences hippocampal function during adolescence. Although the contribution of this pathway remains to be investigated, our findings demonstrate that adolescent light cycle disruption is associated with sex-dependent alterations in hippocampal circadian regulation, plasticity-related signaling, dendritic architecture, and memory. Importantly, the recovery of memory performance and associated molecular alterations following restoration of a standard light-dark cycle suggests that these effects are reversible. Together, these findings highlight the importance of the nighttime light environment for hippocampal function during adolescence and identify potential molecular mechanisms underlying sex-dependent vulnerability to light cycle disruption.

## Materials and methods

### Animal

Female and male mice used in this study CD1 mice. Mice were bred and reared in an on-site animal facility at UofSC under controlled environmental conditions and maintained in standard group housing cages with ad libitum access to normal chow and water. Light intensity was approximately 600 lux at the top of the cage, and the ambient temperature was 23° ± 2°C. The time referred to as “lights on” is defined as ZT0. We minimized the number of animals used in each experiment (sample size based on power analysis) and the animals’ pain and distress. Mouse studies were conducted in accordance with regulations of the Institutional Animal Care and Use Committee at UofSC (Assurance #D16-00028, protocol #2615-101736-090222).

### Chronic Light cycle disruption paradigm

Adolescent mice (post-natal day 30) were transferred into light-tight circadian cabinets (Actimetrics, IL, United States] and subjected to one of the following lighting conditions for a period of 4 weeks: (1) standard 12:12 hr light-dark cycles (12L:12D) (light at ~600 lux) (Control group); (2) 19hr of light and 5hr of darkness for 5 days and 12L:12D for 2 days per week (LCD group); (Figure 1). LED light (~600 lux) is turned on during the night phase of the animals’ cycle, resulting in increased duration of light exposure for 5 days of a 7-day cycle. After 4 weeks of either LCD or control conditions, mice were placed in 12L:12D for 5 days and tested for memory and avoidance behaviors and then sacrificed (Bonilla et al. 2024). For long term exposure, the mice LCD exposed mice were placed back into standard 12:12 hr light-dark cycles (12L:12D) (light at ~600 lux) for 4 weeks. Animals were housed in identical conditions and monitored daily. Light cycle programming and behavioral testing were performed by experimenters blinded to group assignment. After 4 weeks of either LCD or control conditions, mice were placed in 12L:12D for 5 days and tested with novel object recognition or novel object location tasks. For habituation mice were placed in the behavior room for 30 minutes prior behavior testing at ZT8. All behaviors were recorded and analyzed using EthoVision XT (Noldus Information Technology, USA) to track the location of mice.

### Novel Object Recognition (NOR)

The novel object recognition task is performed in 3 consecutive days: On day 1, mice were placed in an open arena (40 cm x 40 cm x 40 cm) and were allowed to explore for 5 minutes before being returned to their home cage. On day 2, mice were placed in the same open arena with two identical quadratic yellow blocks placed on the diagonal at an equal distance from the walls and were allowed to freely explore the objects for 10 minutes. Mice were then returned to their home cage. On day 3, mice were placed in the arena with the familiar object and a novel object which consisted of a quadratic red block and were allowed to explore both objects for 10 min. At the end of each test, the arena and objects were sanitized with 70% v/v ethanol. Mice were recorded using Ethovision software, and the time exploring the familiar and the novel object was calculated as the percentage of time spent with the novel versus the familiar object. The discrimination index (DI) was calculated as the time spent exploring the novel object minus the time spent exploring the familiar object divided by total exploration time. A value below zero describes subjects exploring the familiar object more than the novel object. A value above zero describes animals exploring the novel object more than the familiar object.

### Novel Object Location (NOL)

The novel object location task is performed in 3 consecutive days: On day 1, mice were placed in an open arena (40 cm x 40 cm x 40 cm) and were allowed to explore for 5 minutes before being returned to their home cage. On day 2, mice were placed in the same open arena with two identical quadratic yellow blocks placed adjacent to each other at an equal distance from the walls and were allowed to freely explore the objects for 10 minutes. Mice were then returned to their home cage. On day 3, mice were placed in the arena with the familiar object at the same location and another in a novel location (diagonal to each other) and were allowed to explore both objects for 10 min. At the end of each test, the arena and objects were sanitized with 70% v/v ethanol. Mice were recorded using Ethovision software, and the time exploring the familiar and the novel object was calculated as the percentage of time spent with the novel versus the familiar object. The discrimination index (DI) was calculated as the time spent exploring the novel object minus the time spent exploring the familiar object divided by total exploration time. A value below zero describes subjects exploring the familiar object more than the novel object. A value above zero describes animals exploring the novel object more than the familiar object.

### Immunohistochemistry

After 4 weeks of either LCD or control conditions, mice were placed in 12L:12D for 5 days and then sacrificed at four different ZTs: (in reference to the daily light cycle): ZT2, ZT8 (p-CREB and DBC), ZT16, and ZT22 (all four ZT’s for CRY2). Mice were deeply anesthetized with isoflurane and transcardially perfused with 1% phosphate-buffered saline (PBS) solution immediately followed by 4% paraformaldehyde solution in PBS (pH=7.4) for tissue fixation. Then, the brains were collected, post-fixed in 4% PFA for 24h, cryoprotected in 30% sucrose solution, and stored at 4°C until use. Frozen brains were sectioned (30 μm) with a standard Leica microtome (SM2010R) and stored in cryoprotectant solution at −20°C until use. Six to seven coronal sections from each brain encompassing the hippocampus were processed for immunohistochemistry (IHC) analysis. Immunofluorescent labeling was performed in free-floating slices that were first blocked for 1 hour in 0.1 M phosphate buffer (PB) containing 3% normal horse serum (NHS) and then permeabilized in 0.2% Triton X-100. Slices were then incubated with antibodies: chicken anti-NEUN (1:500; Millipore, ABN91, USA), rabbit anti–p-CREB (1:300; Invitrogen, MA5-11192, USA), guinea pig anti-CRY2 (1:500, Gift from Lamia Lab), rabbit anti-Doublecortin (1:500, Cell Signalling, 4604S, USA) in 1% PBS, 3% NHS, and 0.2% Triton X-100 for 24 or 48 hours at 4°C with constant shaking. After three washes in PBS, slices were incubated for 1 hour at room temperature with fluorescent-tagged secondary antibodies, which were all used at 1:1000 dilution. (1:100; AlexaFluor anti-rabbit 647 nm, AlexaFluor anti-rabbit 555 nm, anti-guinea pig 555 nm, anti-chicken 488). Brain sections were washed, mounted in gelatin on glass slides, counter-stained, and cover-slipped with Dapi Fluoromount-G™ (Electron Microscopy Sciences, USA). Images were taken with a Leica TCS SP8 multiphoton confocal microscopy system (Leica Microsystems, Germany) and LASX software (Leica Microsystems, Germany). Six to three DG sections per animal were used and quantifications were performed blind with ImageJ software. Quantification data were plotted either as the average or as the total number of positive neurons per nucleus per animal.

### RNAscope ® in situ hybridization (ISH)

For each mouse, two slices encompassing the hippocampus were processed for RNAscope in situ hybridization (Multiplex Fluorescent Reagent Kit version 2, Advanced Cell Diagnostics) following the manufacturer’s instructions for detection of Per1 (ACD, catalog #438751), Clock (ACD, catalog #492401), Bdnf (ACD, catalog #424821), and Camkv (ACD, catalog #822941) at 4 different ZTs: ZT2, ZT8, ZT16, and ZT22. Sections were washed in PBS, treated with hydrogen peroxide and mounted in glass slides with 1% PBS. Later, brain sections were incubated in the retrieval buffer and dehydrated in ethanol, previously to a protease treatment. Then, sections were incubated with specific probes for 2 hours at 40°C, and the amplification steps were followed as indicated by the manufacturers. Finally, sections were counter-stained, and cover-slipped with Dapi Fluoromount-G™ (Electron Microscopy Sciences, USA). Images were taken with a Leica TCS SP8 multiphoton confocal microscopy system (Leica Microsystems, Germany) and LASX software (Leica Microsystems, Germany). Quantifications were performed blind with Fiji ImageJ software. Positive area analysis was used to quantify pixels in the specific ROI drawn around the DG (H. Chen et al. 2019).

### Golgi-Cox staining and Dendritic Morphology Analysis

Hippocampal neuronal morphology was assessed using Golgi-Cox staining adapted for cryosection (Zhong et al., 2019). Brains from Control and LCD male and female CD-1 mice sacrificed at ZT8 were extracted, dissected and immersed in Golgi-Cox working solution (5% potassium dichromate, 5% mercuric chloride, and 5% potassium chromate mixed at 5:5:4 with 10 volumes distilled water) for two weeks at room temperature in the dark, with solution renewal at 24 hours. Tissues were then transferred to cryoprotectant solution (20% sucrose, 15% glycerol in distilled water) at 4°C for 48–72 hours until fully sunk. Coronal sections (60 µm) through the dorsal hippocampus were cut on a cryostat (CM1950; Leica Biosystems) at −20°C, mounted onto gelatin-coated slides with 30% sucrose solution, and dried overnight in the dark. Slides were washed in distilled water, developed in 20% ammonia solution for 10 minutes, fixed in 1% sodium thiosulfate for 10 minutes, dehydrated through a graded ethanol series (50– 100%), cleared in xylene, and cover slipped with Permount (Fisher Scientific, SP15-100). Golgi-impregnated neurons in the dentate gyrus were selected for analysis based on complete, untruncated dendritic arbors and clear impregnation free of precipitate artifacts. Dendritic length was quantified in ImageJ (NIH), and dendritic spine density was assessed from (63x) high-magnification images.

### TUNEL Assay

Hippocampal apoptosis was assessed in brain sections from Control and LCD male and female CD-1 mice sacrificed at ZT8 using the Click-iT Plus TUNEL Assay for In Situ Apoptosis Detection with Alexa Fluor 647 dye (catalog no. C10619; Invitrogen, Thermo Fisher Scientific, Waltham, MA) according to the manufacturer’s instructions. Briefly, sections were fixed in 4% paraformaldehyde in PBS for 15 minutes at room temperature, washed twice in PBS, and permeabilized with 0.25% Triton X-100 in PBS for 20 minutes. Sections were then washed and incubated with Proteinase K solution for 15 minutes at room temperature, followed by re-fixation in 4% paraformaldehyde for 5 minutes. After washing, sections were incubated with the TdT reaction cocktail; comprising TdT Reaction Buffer (Component A), EdUTP nucleotide mixture (Component B), and recombinant TdT enzyme (Component C) for 60 minutes at 37°C in a humidified chamber. Sections were washed and incubated with the Click-iT Plus TUNEL Supermix; comprising 1X Click-iT Plus TUNEL Reaction Buffer, Copper Protectant, and Alexa Fluor 647 picolyl azide for 30 minutes at room temperature, protected from light. Neurons were immunolabelled with chicken anti-NEUN (1:500; Millipore, ABN91, USA). As a positive control, parallel sections were treated with DNase I (catalog no. 18068-015; Invitrogen) for 30 minutes at room temperature prior to the TdT reaction to induce DNA strand breaks. TUNEL-positive (apoptotic) cells in the dentate gyrus (two sections per animal) were imaged by fluorescence microscopy and quantified across matched hippocampal sections per animal.

### Statistical Analysis

Investigators who participated in end point analyses were blinded to the light protocol. Statistical analysis was carried out using GraphPad Prism (GraphPad Software, La Jolla, CA, USA). For normally distributed data, a parametric test was used [one-way or two-way analysis of variance (ANOVA) or Student’s t-test and p-value less than 0.05 was considered significant and was indicated as followed: * p<0.05; ** p<0.01; *** p<0.001; **** p<0.0001. The statistical tests performed for each experiment are indicated in the figure legends.

## Acknowledgements

We thank all members of the Porcu’ Laboratory for valuable discussions and comments. This work was supported by NIH-NCCIH grant #AT010903 and #AT013234 to A.P, and SPARC grant (#80007606) to A.B.

